# SortIT — A Tool For Assessing Observer Variability And Creating Ground Truth Image Classification Datasets

**DOI:** 10.64898/2026.05.28.728616

**Authors:** Wataru Uegami, Tom Bisson, Ethan N. Okoshi, Frederico Gaia Costa da Silva, Jijgee Munkhdelger, Kris Lami, Norman Zerbe, Andrey Bychkov, Junya Fukuoka

## Abstract

Interobserver variability in pathological assessments is a well-recognized challenge that impacts diagnostic reliability and disease understanding. This variability exists across many subspecialties due to the subjective nature of evaluations. Artificial intelligence (AI) applied to whole slide images has potential to standardize procedures and reduce variability in pathology, but transitioning to these technologies does not guarantee improvement. Establishing reliable ground truth datasets with consensus annotations is crucial for developing robust AI solutions. We introduce SortIT, an open-source web application that facilitates systematic creation and evaluation of ground truth image tile annotations. SortIT enables multiple annotators to independently label tiles, with flexible user permission controls. Annotated data can be exported for statistical analysis of observer variation and for creating ground truth datasets from consensus tiles. We outline protocols using SortIT for several use cases: (1) mitosis segmentation in tumor regions, (2) evaluating AI solutions for prostate cancer grading by comparing to expert consensus, and (3) granuloma classification by annotating discriminative tile-level features. Key strengths of SortIT lies in its ease of deployment, making it accessible and usable for a wide range of users. Overall, SortIT provides a valuable tool to establish high-quality ground truth datasets and comprehensively assess observer variability. Critical evaluation of ground truth quality using systematic annotation methodologies is crucial for developing accurate and generalizable diagnostic AI tools. Its open-source nature facilitates community adoption and further development.

## 1 Introduction

In pathology, interobserver variability in tissue evaluations is a common yet critical issue. It not only affects diagnostic reliability at the clinical level but also hinders the understanding of diseases. As pathological assessment often relies on the subjective judgment of pathologists, this problem arises in nearly every subspecialty and domain of pathology, including disease classification, tumor grading, cancer staging, and various reporting parameters. There are many notable examples of considerable expert disagreement in breast,^1^ prostate,^2^ lung,^3^ skin,^4^ gynecologic,^5^ and endocrine tumor^6, 7^ pathology, among others. The same issue is equally recognized in non-neoplastic diseases^8–10^and cytopathology.^11^ Furthermore, evaluation of common ancillary techniques, such as immunostaining and in situ hybridization, have also been shown to be subject to interobserver variability.^12, 13^ We conducted a survey of international expert pathologists (*n* = 81; December 2025), distributed via social media, to assess their estimation of interobserver variability across different diagnostic domains (Figure 1). The results showed that pathologists believe agreement is generally high for common neoplastic and non-neoplastic diseases, while uncertainty and discordance increase when rare entities are encountered. For malignant vs. benign classification of common neoplasms, 84% (95%CI [0.74, 0.91]) of respondents believed that interobserver agreement was ≥80%, compared with 59% (95%CI [0.48, 0.70]) for rare neoplasms.

**Figure 1.**
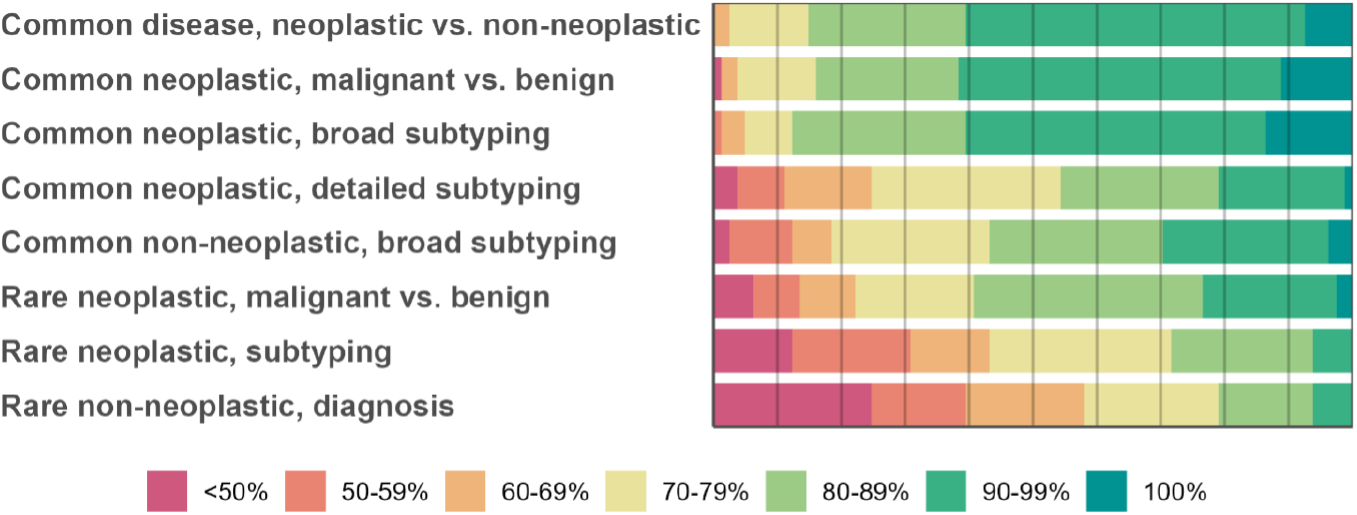
Survey results from expert pathologists evaluating the level of interobserver agreement on various tasks. The survey was shared via social media, and 81 responses were collected over 2 weeks in December 2025. Each question was multiple choice, and only one answer could be selected.

Using whole slide images (WSIs) analyzed with computational pathology techniques such as artificial intelligence (AI) could help standardize diagnostic procedures and potentially reduce interobserver variability, thus optimizing patient care.^14^ Validation studies comparing digitized WSI-based workflows with conventional microscopy have shown mixed results. Some studies, on renal pathology^15, 16^ and on tumor bud evaluation,^17^ showed improved agreement with WSI, while others have shown better agreement using conventional microscopy.^18, 19^ AI assistance has been shown to improve interobserver agreement on WSI in several studies.^20, 21^ However, the use of AI in pathology carries the risk of overinterpreting the performance metrics achieved with these computational tools. When AI models achieve high performance metrics, there is a tendency to develop a certain level of trust in the algorithm. It is important to note that these metrics are based on the algorithm’s performance on a pre-existing validation or test data set. Yet, the quality and representativeness of this specific data set is rarely critically examined. This raises important considerations regarding interobserver variability and how it might be counterbalanced, strategies to deal with tissue structures lacking consensus between the rating pathologists, possible distinctions between structures with immediate consensus and those necessitating discussion, or potential diagnostic variations across different laboratories. These and other aspects of dataset compilation must be considered when applying computational pathology tools to routine diagnostics. To deal with this fundamental issue, a variety of approaches have been developed to measure and, when necessary, minimize the effects of interobserver variability.^22^ In this paper, we present an image sorting tool for dataset building, SortIT. SortIT is a web application that allows users to compile image datasets with human-confirmed ground truth labels. Our patch-level labeling tool is engineered for ease of use, offering an intuitive and user-friendly experience. SortIT can be deployed locally or in the cloud, ensuring easy access. The tool includes a user management system which maintains a clear differentiation between administrators and regular users. Importantly, it allows for granular permission assignments to individual users, enabling tailored setups that align with the specific needs of various institutions. SortIT facilitates independent labeling by multiple users, offering a high degree of flexibility. SortIT has been made available as open-source software (https://github.com/FukuokaLab/SortIT).

## 2 Methods

### 2.1 Brief Overview

The workflow begins with collecting a set of WSIs on which tile-level classification should be performed. Then, the image tiles need to be generated from these WSIs in order to import them into SortIT. Subsequently, several pathologists can perform the classification task via the browser-based user interface. When the classification phase is completed, the resulting data can be exported as a CSV file, facilitating a thorough evaluation and comprehensive analysis (Figure 2). Usage of the tool is described in detail in the code repository README.

**Figure 2.**
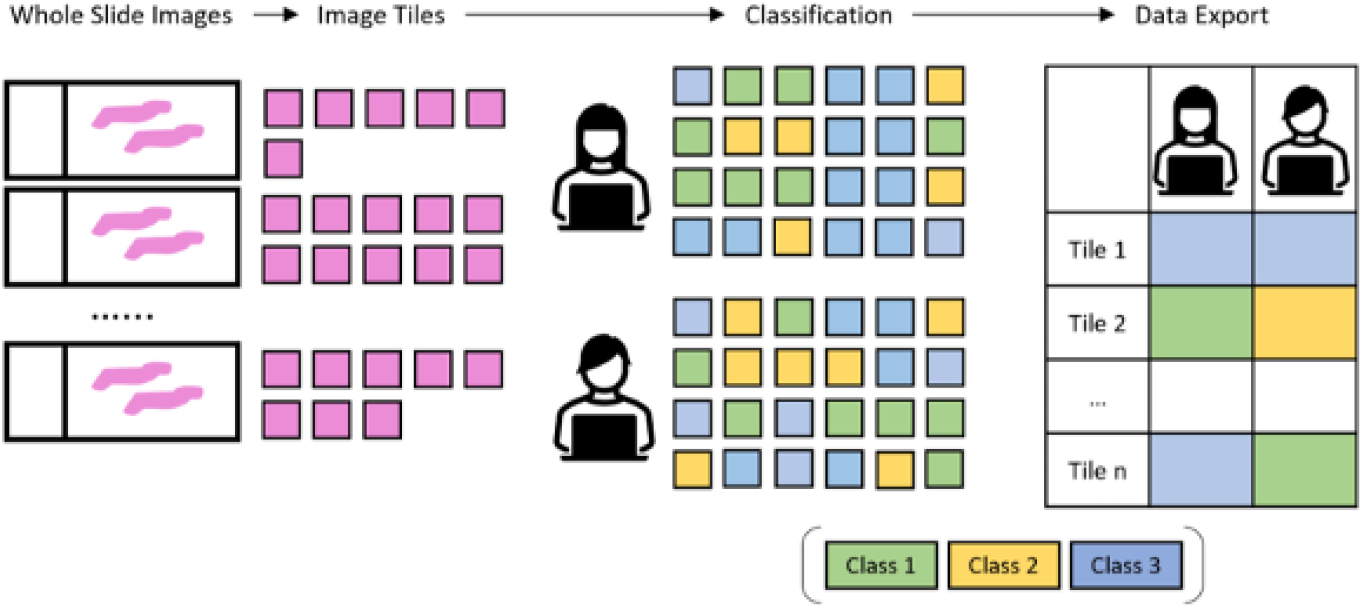
Visualization of the complete workflow. Image tiles need to be generated from a set of WSIs to then be classified by multiple pathologists. Afterwards, the results can be exported as a CSV file for further analysis.

### 2.2 Software

SortIT is a web application using the Django framework. It can be accessed from any device which can support a web browser, including smartphones, tablets, and personal computers. Within the software, user roles are divided into administrators and standard users. Administrators have additional privileges, such as preparing datasets and managing user accounts. By default, standard users are limited to image annotation, although administrators can expand their access as needed. The annotation process can follow either a single-class or a multi-class approach. In the single-class annotation mode, images are displayed in a grid format, allowing users to select images which do not belong to the target class by clicking on the images (Figure 3a). Selected images are filtered out of the dataset (label is left empty). In the multi-class mode, images are shown individually, and users select from multiple applicable class options for each image (Figure 3b). A progress indicator informs users how many images have been annotated out of the total batch, helping monitor workflow completion. A key feature of SortIT is its ability to export annotated data in CSV format. In this format, users are represented by rows, images by columns, and the assigned classes are recorded in the corresponding cells. This exported CSV file offers a structured summary of the annotations, making it suitable for further analysis.

**Figure 3.**
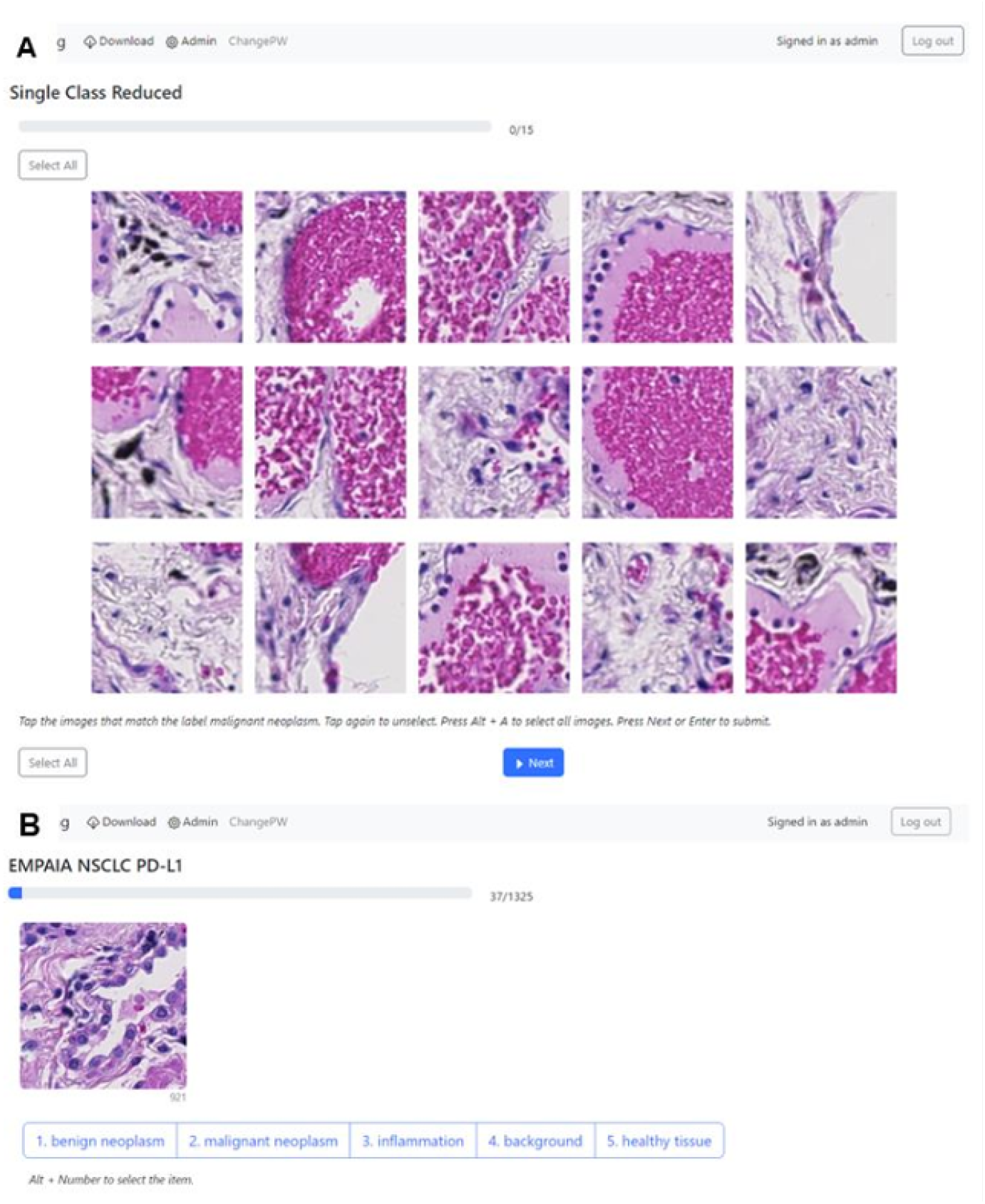
SortIT user interface. Single-class (A) and multi-class (B) sorting approaches.

### 2.3 Tile Creation

The presented software is based on classification at the image tile level, with no special requirements for the sizes of the image tiles. These may already correspond exactly to the dimensions expected by the neural network used in each case, but the tool does not restrict in this regard, so that different approaches to ground truth creation can also be applied. Generating these image tiles is not part of SortIT. The authors generally use openslide-python or QuPath. For this purpose, the QuPath documentation provides a sample script that can be used to configure various parameters such as tile size, size of the overlapping area, resolution, or consideration of any annotations that may be present. In some scenarios it is advantageous to modify the image tiles afterwards. One possibility, for example, is to display a larger tissue area and additionally mark the area to be classified. This could ensure that the surrounding tissue can also be considered during the sorting process. Furthermore, users can apply methods such as MIXTURE to further reduce the burden of labeling images.^23^

### 2.4 Statistical Evaluation

Ground truth quality assessment requires precise statistical analysis of interobserver and/or intraobserver variability. This is typically achieved through interobserver agreement measures such as Cohen’s kappa and Fleiss’ kappa. Our tool facilitates these analyses by enabling CSV data export. To determine the interobserver variability, it is sufficient to create an individual user account for each annotating expert. Depending on the problem under investigation, it is advisable to define certain procedures. For example, a maximum session duration could be defined to minimize fatigue. Other problem-specific study parameters could be, for example, compliance with guidelines, defining classes by the presence of morphological characteristics, or the use of cutoff-values. To determine intraobserver variability, several user accounts could be created for a single person, with all of these accounts operating on the same data set. First, this data set should be fully annotated with one account. Then, a washout period should be applied, usually of two weeks or longer. Subsequently, the dataset can be annotated by the same pathologist again using a separate account.

### 2.5 Customizing SortIT

SortIT is built using the Django^24^ framework. New features can be added to the app by altering the code found in our public repository. Adding new features will require familiarity with Python and web design languages like HTML and JavaScript, but the website is designed to be simple to extend. Detailed discussion of the codebase and its structure is included in the README file in the code repository.

## 3 Results

### 3.1 Validation Study

To validate SortIT with an empirical study, three board-certified pathologists (P1–P3), who were blinded to the study hypothesis and comparative objectives, independently performed tumor-rich patch selection on two cases. Whole-slide images were pre-segmented into non-overlapping 224 *×* 224 pixel patches. Using the SortIT interface, each pathologist selected patches estimated to contain ≥70% tumor cells based on visual assessment. For comparison, manual annotation was performed using QuPath (version 6.0.0). Tumor regions were delineated at the structural/glandular level using polygon annotations. Subsequently, 224 *×* 224 pixel patches were generated from areas in which ≥70% of the patch area was covered by the annotated tumor region. The number of tumor-rich patches identified by each method and the time required for completion were recorded for each pathologist and case. SortIT yielded significantly higher numbers of tumor-rich (≥ 70%) image acquisitions compared with manual annotation (paired t-test, *p* = 0.0074, 95%CI=[106.1, 414.2]; Wilcoxon signed-rank test, *p* = 0.031, 95%CI=[105, 531]). There was no statistically significant difference in acquisition time in seconds between SortIT and manual annotation (paired t-test, *p* = 0.50, 95%CI=[-384.81, 687.48]; Wilcoxon signed-rank test, *p* = 0.44, 95%CI=[-647, 900]). (Figure 4)

**Figure 4.**
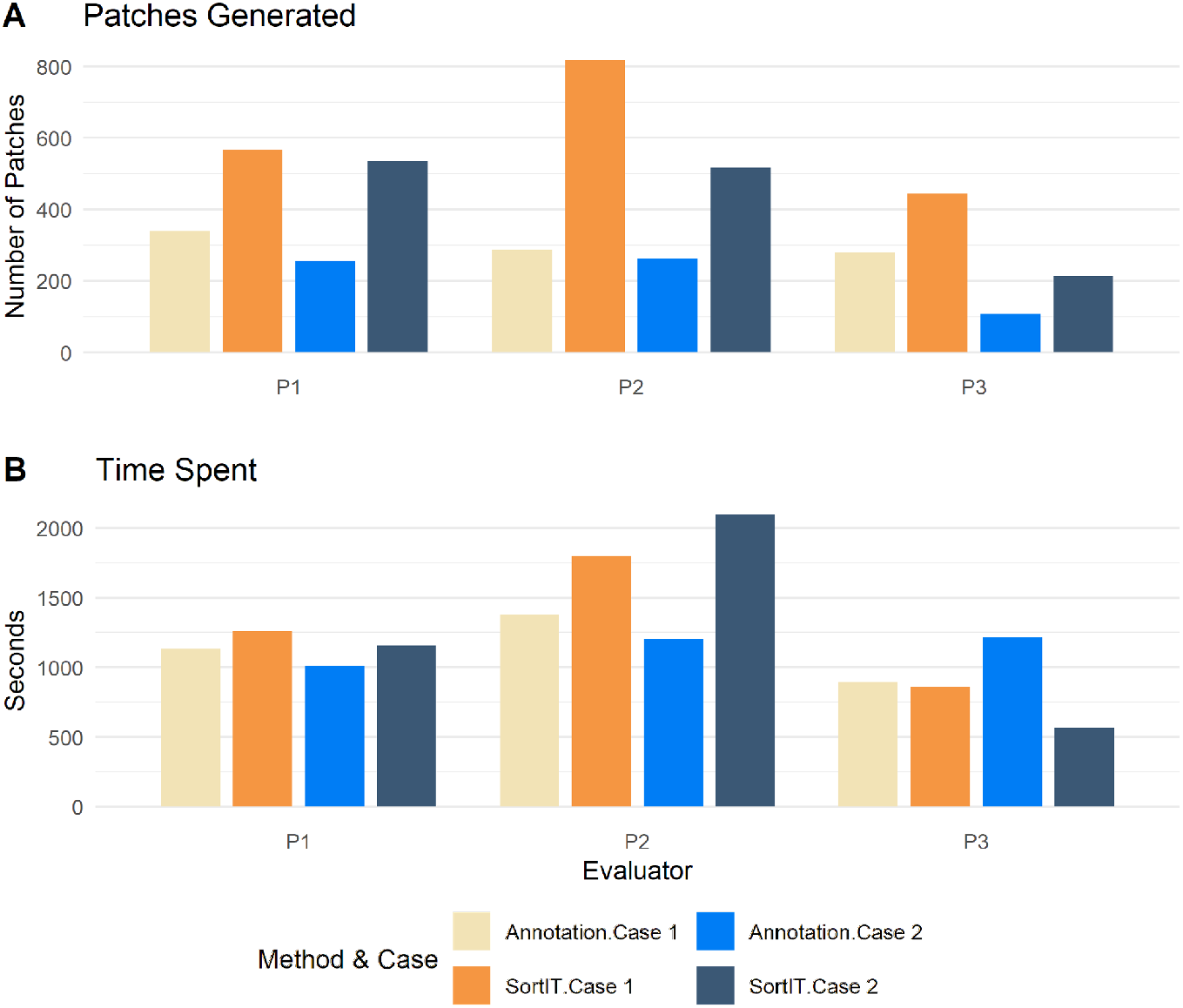
Validation study results between hand annotations vs. SortIT for patch dataset creation. 2 cases were evaluated by 3 trained pathologists. (A) Number of patches generated by each pathologist. (B) time spent on each task.

### 3.2 Use Cases

SortIT provides a useful tool to establish ground truth datasets composed of consensus tiles^3^ and comprehensively assess interobserver and intraobserver variability. To facilitate these evaluations, we designed different study protocols which can serve as guidelines, ensuring a systematic approach to studying and quantifying the levels of variability within our analyses. In the following, we will delineate several distinct methodologies for establishing ground truth for various tasks using SortIT.

#### 3.2.1 Use Case 1: Mitosis Segmentation

In this first use case we describe how we would train a neural network for mitosis segmentation in H&E images. As input, we would define a region of interest (ROI) within the WSI, outlining tumor areas exclusively to accurately count relevant mitotic figures. As there is no need to address inherent spatial context between individual mitoses, the AI will work solely at the tile level. By building a dataset comprising the precise coordinates of mitoses, we gain the flexibility to generate image tiles containing mitoses positioned at various locations, scales and rotations. As there might be more than just one mitosis in one image tile, we will aim for a segmentation model to find all instances of mitosis in each specific tile. To generate the image tiles to be classified, we employed the QuPath cell detection algorithm to achieve automated segmentation of cells within WSI. It results in a set of annotations outlining both the outer cell membrane as well as the nuclei. Leveraging this annotation data, we proceeded to create tiles cropped to the dimensions of the detected cells using a custom groovy script.^25^ In this case, there is no need for a specific tile size since the training data is generated at a later stage utilizing the output of SortIT. When all cells were classified into either mitotic or non-mitotic in single-class approach, we can map the IDs of the images to the results obtained from QuPath’s cell detection algorithm resulting in a comprehensive dataset, featuring segmentation masks for both mitotic as well as non-mitotic cells. The non-segmented remainder of the ROI may serve as a background class allowing a 3-class segmentation model to be trained (Figure 5).

**Figure 5.**
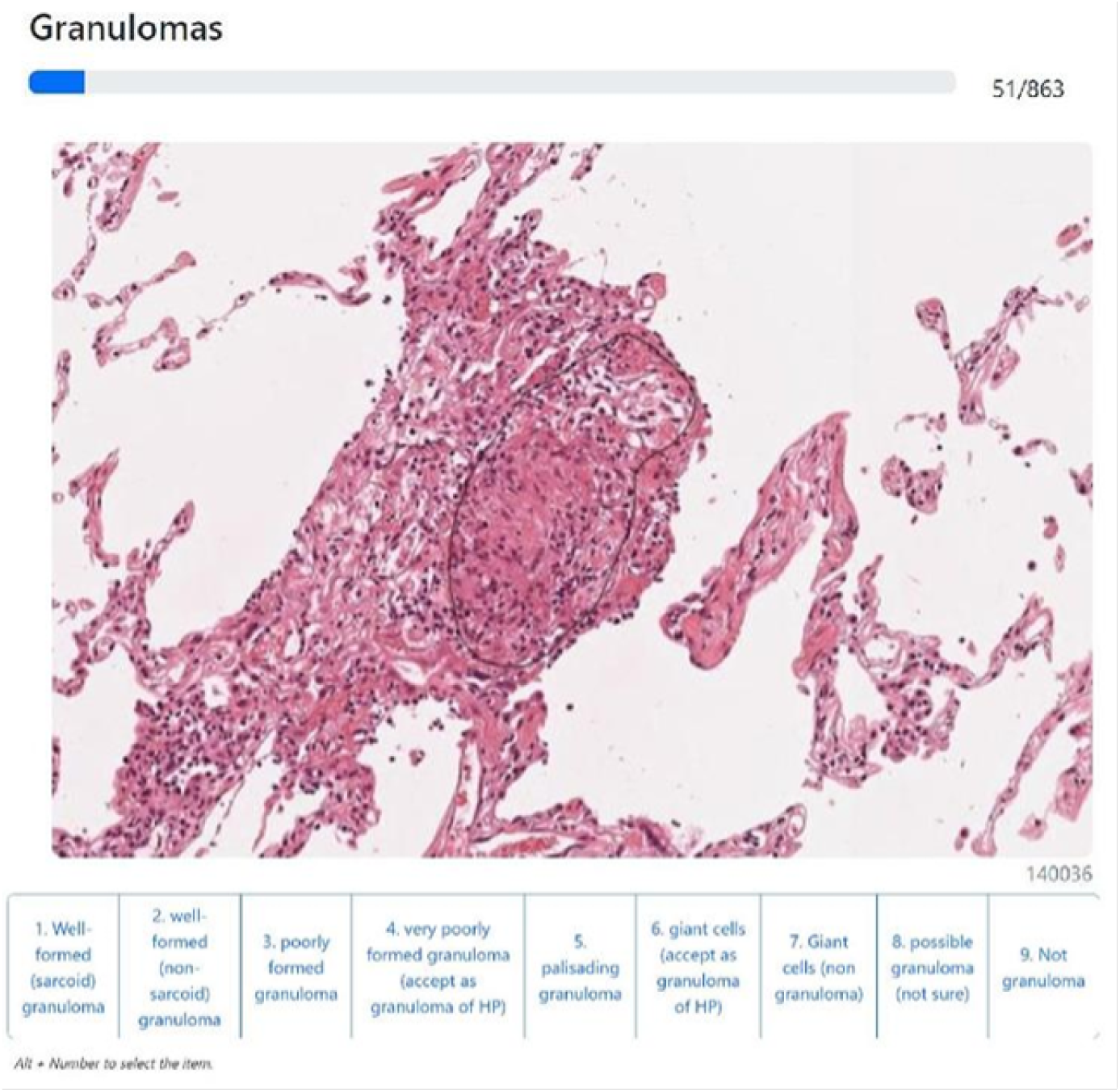
Overview of dataset creation for ground truth generation in mitosis segmentation.

In evaluating the ground truth for mitosis detection in H&E slides, it’s important to acknowledge the inherent challenge of achieving perfect classification accuracy for pathologists. Given this, we recommend creating a representative subset, encompassing a percentage of the entire dataset, for annotation by pathologists allowing for a comprehensive interrater agreement assessment using Fleiss’ kappa. By deriving insights from their annotations, we obtained valuable information regarding the dataset’s reliability and the challenging nature of the task. Understanding the imperfections in manual annotation of mitosis, we are confident that with a sufficiently large dataset, the neural network will effectively mitigate the noise, encompassing both falsely annotated cells and potentially missed mitotic figures, within the dataset.

#### 3.2.2 Use Case 2: Evaluation of AI-Solutions for Prostate Cancer Grading

In the second use case, we demonstrate how to analyze the performance of AI tools for prostate cancer grading. The relevance of this topic is reflected in the continued emergence of commercial AI solutions in this field, such as those from Ibex, Paige Prostate or Ibex Galen prostate, to name a few. To investigate these solutions, they are first employed on a series of prostate slides to then evaluate the individually identified lesions. In doing so, both the AI-assigned Gleason grades and the accuracy of the annotations will be examined. To obtain the individual images, the respective AI algorithm to be evaluated is applied to the WSIs followed by acquiring screenshots of each lesion either directly through the program’s user interface or alternatively in QuPath and saving them with the slide name, a unique ID for the lesion, and the Gleason grade provided by the tool. Then, all images are uploaded to SortIT and made available to multiple pathologists for evaluation (Figure 6).

**Figure 6.**
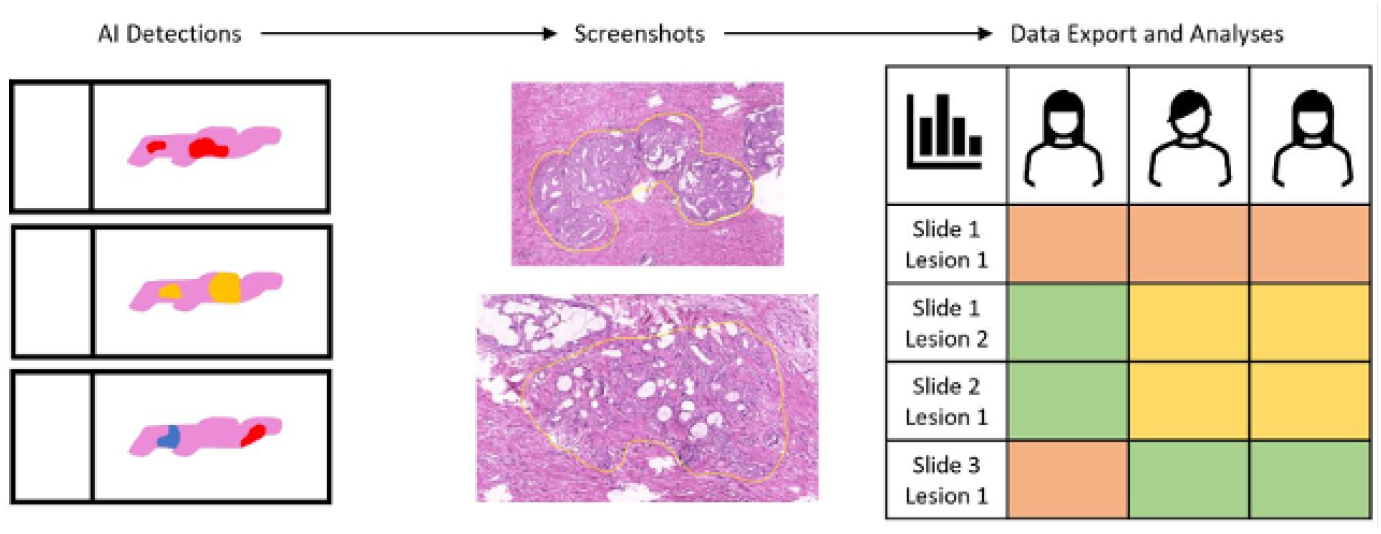
Overview of the AI evaluation process for prostate cancer grading.

For the evaluation, Cohen’s kappa is calculated to measure the agreement between the AI tool and each pathologist in grading the lesions. In addition, Fleiss’ kappa is calculated between all pathologists for each lesion, providing information on the overall agreement between pathologists in grading. Furthermore, the percentage of lesions for which the AI-assigned Gleason grade is in agreement with the majority pathologists is determined, illustrating the agreement of the prostate AI tool with expert opinion. Finally, the percentage of cases in which the AI yields false positives – defined as giving a grade differing from the majority consensus – is determined, providing insight into its specificity. False negatives are not included in this process but can also be covered by adjusting data generation accordingly. This comprehensive investigation ensures a solid understanding of the performance of the AI tool and helps assess its applicability in the diagnosis of prostate cancer.

#### 3.2.3 Use case 3: Granuloma classification

Granulomas, organized aggregates of immune cells, form as a defense mechanism in response to persistent infections, foreign materials, or autoimmune conditions. In the lungs, granulomas are commonly caused by diseases such as tuberculosis, non-tuberculous mycobacterial infections, sarcoidosis, hypersensitivity pneumonitis, and granulomatous vasculitis. Each of these diseases have distinct histological features in their granulomas, but they do not always have clear boundaries and often show overlapping characteristics. Each of these diseases displays unique histological features in their granulomas, 6but overlapping characteristics often make differentiation difficult, complicating the diagnostic process. Accurately identifying the underlying disease, however, is crucial for delivering effective therapy. Necrotizing, well-formed granulomas are associated with infections, well-formed granulomas with fibrosis are seen in sarcoidosis, small and poorly formed granulomas composed of loosely connected histiocytes are characteristic of hypersensitivity pneumonitis, and well-formed granulomas with a palisading pattern are observed in granulomatosis with polyangiitis and other types of vasculitis. These features are not necessarily uniform across the entire granuloma. Consequently, certain regions may reflect the underlying disease more accurately than others. AI algorithms are designed to optimize their decision-making over all training data points, necessitating robust and precise annotation strategies to create a reliable ground truth. When regarding granuloma, it might seem intuitive to assign a single label to an entire granuloma, describing the distinct characteristic reflecting the underlying disease. However, because the specific histological features of certain granulomas can be concentrated in localized areas, there is a risk that a neural network will optimize towards irrelevant or non-disease-specific features across the whole structure. To address this, we use SortIT to annotate granulomas at the tile level using a multi-class approach, increasing the likelihood that AI will accurately learn the characteristic histological features (Figure 7).

**Figure 7.**
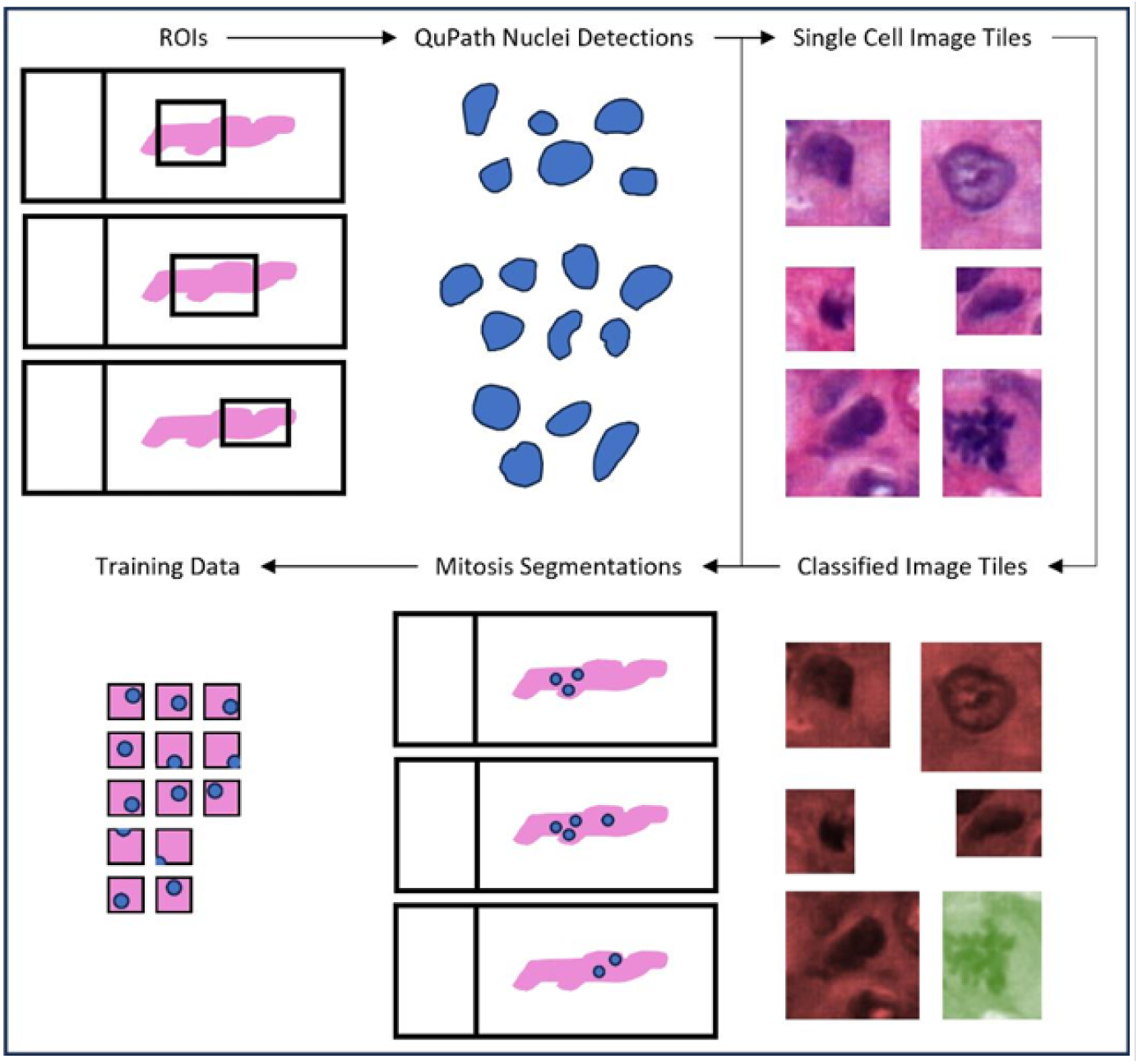
Overview of the AI evaluation process for prostate cancer grading.

## 4 Discussion

The developed SortIT tool is an open-source, web-based application designed for the labeling of image tiles. Its key advantage lies in its ease of deployment, making it accessible and usable for a wide range of users. The versatility of the tool allows it to serve various use cases in the field of digital pathology, ranging from the pure investigation of intra- and interobserver variabilities up to the creation of a ground truth used to train an AI and simultaneously measure the quality of the dataset. Since the respective evaluations are performed using data that has been collected and processed automatically, not only is the labeling process simplified, but also, the tool facilitates evaluation of the reliability and precision of the collected data. There are a number of available tools for the purpose of labeling image data, including CVAT, VoTT, MakeSense, Label Studio, OpenLabeler, MedTagger, etc. However, many of these tools are no longer actively maintained, are not open source, focus more on object detection and segmentation tasks, or are more complex. We designed SortIT to be limited in scope so as to be as simple as possible to use for the specific use case. The main feature of SortIT, the ability for multiple users to enter annotations for images, is not a common feature among the labeling tools we found. MedTagger and LOST also feature collaborative labeling functionality, but the former is no longer maintained as of 2020, and the latter has a much wider scope. SortIT provides a tool for users that value a fast and simple tool for image classification tasks. One strength of our solution is that it operates as a web application, accessible through a browser, without the need for an installation process. This design not only improves usability but also allows pathologists to conduct data analysis and creation without extensive time, effort, and IT expertise. SortIT can be encapsulated in a Docker container, i.e. a virtual containerized operating system with comprehensive system setup. This not only simplifies the initial setup of the tool but also ensures that the tool can be used regardless of the system environment. Additionally, the tool was implemented using the Django web framework, which is written for Python. Python is widely used in the AI community due to its numerous existing libraries for AI and image processing, such as PyTorch, TensorFlow, Scikit-Image and OpenCV. This, along with the open-source license under which the tool is provided, opens up multiple opportunities for further development by the community. One of the main hurdles for the development, acceptance and dissemination of AI solutions is the availability of annotations. While there are a number of publicly available datasets,^26, 27^ this is not true for every single diagnostic domain. Not every research group is sufficiently staffed and funded to develop their own annotation tools and processes. Thus, there is a demand for making annotations more widely available.^28^ SortIT counteracts this by greatly simplifying the annotation/sorting process. Another significant benefit is the lack of having to transfer data between individual annotators, allowing seamless collaboration across multiple sites. Since SortIT is provided as open source, anyone can create another ground truth using exactly the same annotation methodology and can furthermore measure and compare the inherent interobserver variability in these datasets. A notable constraint of this tool is its limitation to tile-level classification tasks, constricting its versatility and applicability for image segmentation tasks. Moreover, in its current form, SortIT does not provide a comprehensive end-to-end solution encompassing the entire process from generating image tiles from WSI to statistically evaluating results. The responsibility to generate image tiles falls on the user, although guidance and a script for a specific case within QuPath are provided to ease this process. Similarly, the tool leaves the statistical evaluation of results to the user, exporting only raw classification data. It is crucial to underline that the evaluation results heavily hinge on the chosen annotation methodology. The current methodology involves classification based on predefined image tiles (still images) without the ability to navigate within the tissue offered by WSI, meaning users are unable to view the surrounding tissue of the tile to put it into context. However, users can choose to upload tiles at any magnification and resolution, so they can ensure that tiles have sufficient context for their specific labeling task.

## 5 Conclusion

The quality of ground truth in a dataset is a critical factor in the development of AI-powered diagnostic solutions and is determined not only by the number of data points included, but also by a transparent explanation of the creation process. Our SortIT tool provides researchers with the ability to perform such an assessment with minimal setup effort. We present several use cases, including classification at the individual cell level, scoring within fixed size areas, as well as tissue structures within their spatial context. It is of great importance that researchers critically evaluate their ground truth by performing such comprehensive quality measurements.

## Notes

https://github.com/FukuokaLab/SortIT

### Competing Interest Statement

The authors have declared no competing interest.

### Summary of Updates

Author list correction. (Jijgee Munkhdelger and Kris Lami are added. One author removed)

